# Targeting and anchoring the mechanosensitive ion channel Piezo to facilitate its inhibition of axon regeneration

**DOI:** 10.1101/2024.12.13.628292

**Authors:** Qin Wang, Leanne Miles, Shuo Wang, Harun N. Noristani, Ernest J. Monahan Vargas, Jackson Powell, Sean J O’Rourke-Ibach, Shuxin Li, Yuanquan Song

**Affiliations:** Raymond G. Perelman Center for Cellular and Molecular Therapeutics, The Children’s Hospital of Philadelphia, Philadelphia, PA 19104, USA; Department of Pathology and Laboratory Medicine, University of Pennsylvania, Philadelphia, PA 19104, USA; Shriners Hospitals Pediatric Research Center (Center for Neurorehabilitation and Neural Repair), Lewis Katz School of Medicine at Temple University, Philadelphia, PA 19140, USA; Department of Neural Sciences, Lewis Katz School of Medicine at Temple University, Philadelphia, PA 19140, USA

## Abstract

Mechanical force orchestrates a myriad of cellular events including inhibition of axon regeneration, by locally activating the mechanosensitive ion channel Piezo enriched at the injured axon tip. However, the cellular mechanics underlying Piezo localization and function remains poorly characterized. We show that the RNA repair/splicing enzyme Rtca acts upstream of Piezo to modulate its expression and transport/targeting to the plasma membrane via Rab10 GTPase, whose expression also relies on Rtca. Loss or gain of function of Rab10 promotes or impedes *Drosophila* sensory neuron axon regeneration, respectively. Rab10 mediates the cell surface expression of integrin β1 (Itgb1)/mys, which colocalizes and genetically interacts with Piezo, facilitating its anchorage and engagement with the microenvironment, and subsequent activation of mechanotransduction to inhibit regeneration. Importantly, loss of Rtca, Rab10 or Itgb1 promotes CNS axon regeneration after spinal cord injury or optic nerve crush in adult mice, indicating the evolutionary conservation of the machinery.

## Introduction

Neurons have varying capacities to regenerate, and nerves in the mature mammalian central nervous system (CNS) retain lower regenerative potential than those in the peripheral nervous system (PNS)^1,2^. Intrinsic regeneration potential varies across types of neurons, including subtypes of retinal ganglion cells (RGCs) and dorsal root ganglion (DRG) neurons^3,4^. Therefore, it is crucial to identify genetic programs that regulate regeneration to provide effective treatments for patients who live with the debilitating effects of nerve damage, ranging from traumatic brain and spinal cord injuries to blindness. Numerous efforts have been made to characterize essential genes and pathways that either promote or inhibit regeneration in the hopes of identifying key therapeutic targets, but current therapies lead to mild recovery at best^5,6^.

Utilizing a *Drosophila melanogaster* neural injury paradigm, we have shown that fly sensory neurons exhibit class-specific regeneration capabilities that power the screening of pro- and anti-regeneration factors^7^. In particular, the RNA repair/splicing pathway has emerged as an inhibitory machinery for axon regeneration. The removal of Rtca, an RNA 3’-terminal phosphate cyclase and the core component of this pathway, improved axon repair in the CNS and PNS, while its overexpression suppressed axon regrowth^8^. Whereas *Xbp1* mRNA splicing and the unfolded protein response (UPR) are the key steps modified by Rtca^8,9^, the events regulating axon regrowth downstream of Rtca remain sparsely studied.

The effect of mechanical forces on neuronal growth has long been observed. However, little was known about the underlying molecular and cellular machinery. Our prior work showed that Piezo inhibits axon regeneration via its function as a mechanosensitive ion channel. Upon sensing the cellular mechanical force during regeneration, Piezo triggers calcium influx in the growth cone and activates the CamKII-Nos-Atr pathway in fly and mammalian neurons^10,11^. Mutant Piezo protein with prolonged channel opening suppresses axon regeneration, while another mutant that cannot conduct ions fails to inhibit regrowth^11^. However, the million-dollar question remained unanswered – what controls the expression, localization and activation of Piezo channels? How are Piezo channels mobilized and gated? Understanding and utilizing the mechanotransduction pathway may offer a new dimension to tackle challenges in neural repair and beyond.

Here, we uncovered genetic and cellular mechanisms underlying Piezo localization and function during regeneration. Upstream, the expression of Piezo is modulated by Rtca, which mediates the expression of both Piezo and Rab10, a small Rho GTPase known to function in endosome transport and recycling^12^. Rtca mediates proper membrane localization of Piezo channels via Rab10, which is enriched at the injured axon tip. Loss of Rab10 promotes while its overexpression impedes axon regeneration in fly sensory neurons. Furthermore, Rab10 mediates the cell surface expression of integrin β1 (Itgb1)/mys, which facilitates Piezo’s anchorage and subsequent activation of mechanotransduction to inhibit axon regeneration. Itgb1/mys loss of function (LoF) drastically enhances fly sensory neuron axon regeneration. Importantly, loss of Rtca, Rab10 or Itgb1 boosts CNS axon regeneration in adult mouse spinal cord injury or optic nerve crush models, indicating evolutionary conservation. Our work thus reveals a multi-step molecular and cellular machinery controlling Piezo function, filling the knowledge gap in the coordination of various regeneration inhibitors, and generates a network of regeneration brakes with hierarchy – the targeting of which provides promising opportunities for therapeutic intervention after nerve damage.

## Results

### The RNA repair/splicing pathway regulates axon regeneration

The cellular stress sensor *Xbp1* is known to undergo non-conventional mRNA splicing, mediated by the RNA repair/splicing pathway^13,14^. Properly spliced *Xbp1* mRNA is required for resolving ER stress, as experienced by injured neurons^8^. *Xbp1* splicing requires the ligase RtcB (RNA 2’,3’-cyclic phosphate and 5’-OH ligase), whose enzyme activity is boosted by the catalyst Archease^14^. As a counteraction, Rtca slows down *Xbp1* splicing. Our prior work has demonstrated that LoF of Rtca or Archease promotes or impedes axon regeneration, respectively^8^. The role of RtcB, however, remains enigmatic. In *C. elegans*, RtcB has been reported to inhibit axon regeneration, independent of the unfolded protein response and Archease^15^. This seeming discrepancy brings up a key question – does RtcB have a conserved role in regeneration or rather function in a species-dependent manner? To address this issue, we generated *RtcB* LoF mutants – *RtcB*^1–^*^2C^*and *RtcB*^1–^*^4B^* by using CRISPR to target a region around 150 bp from the start codon. This results in a 4 and 5 bp deletion of the coding sequence, respectively, predicted to cause frameshift and protein truncation (Figure 1A). To assess axon regeneration, we used class IV dendritic arborization (C4da) neurons (labeled by *ppk-CD4tdGFP*), which robustly regenerate in wildtype (WT) after laser axotomy in the PNS, and found that *RtcB*^1–^*^4B^* showed decreased regeneration capacity (Figures 1B and 1C). Next, we examined transheterozygotes of *RtcB*^1–^*^2C^/Df(2L)Exel8041* to avoid the potential off-target effect of gRNAs. *Df(2L)Exel8041* contains a deletion of a stretch of the chromosome that covers the *RtcB* locus. RtcB LoF significantly reduced the percent of regenerating C4da neurons and the regeneration index, which represents normalized axon regeneration length (Figures 1B, 1C and Methods). The phenotype is comparable to that of the *Archease* LoF mutant^8^, *Archease^PBc^*^01013^ (*Archease^PB^*). Moreover, we found that compared with single heterozygotes of *RtcB*^1–^*^4B^*or *Archease^PB^*, in which C4da neurons exhibit normal regeneration ability, *RtcB* and *Archease* transheterozygotes (*RtcB*^1–^*^4B^/+; Archease^PB^/+*) showed substantially reduced regeneration capacity (Figures 1D and 1E), supporting the hypothesis that RtcB and its cofactor Archease cooperate to allow *Xbp1* splicing and hence axon regeneration (Figure 1A). We then wondered if Archease itself is capable of conferring axon regeneration in neurons that normally fail to regrow. We turned to class III dendritic arborization (C3da) neurons (labeled by *19-12-Gal4>CD4tdGFP; repo-Gal80, nompC-QF*>*mtdTomato* or *nompC-QF*>*mCD8GFP*), which regenerate poorly in WT even after PNS injury. While RtcB overexpression was sufficient to promote axon regrowth, C3da neuron-specific overexpression of Archease was unable to enhance their regeneration (S1 Figure), suggesting that the function of RtcB may be rate limiting.

**Figure 1.**
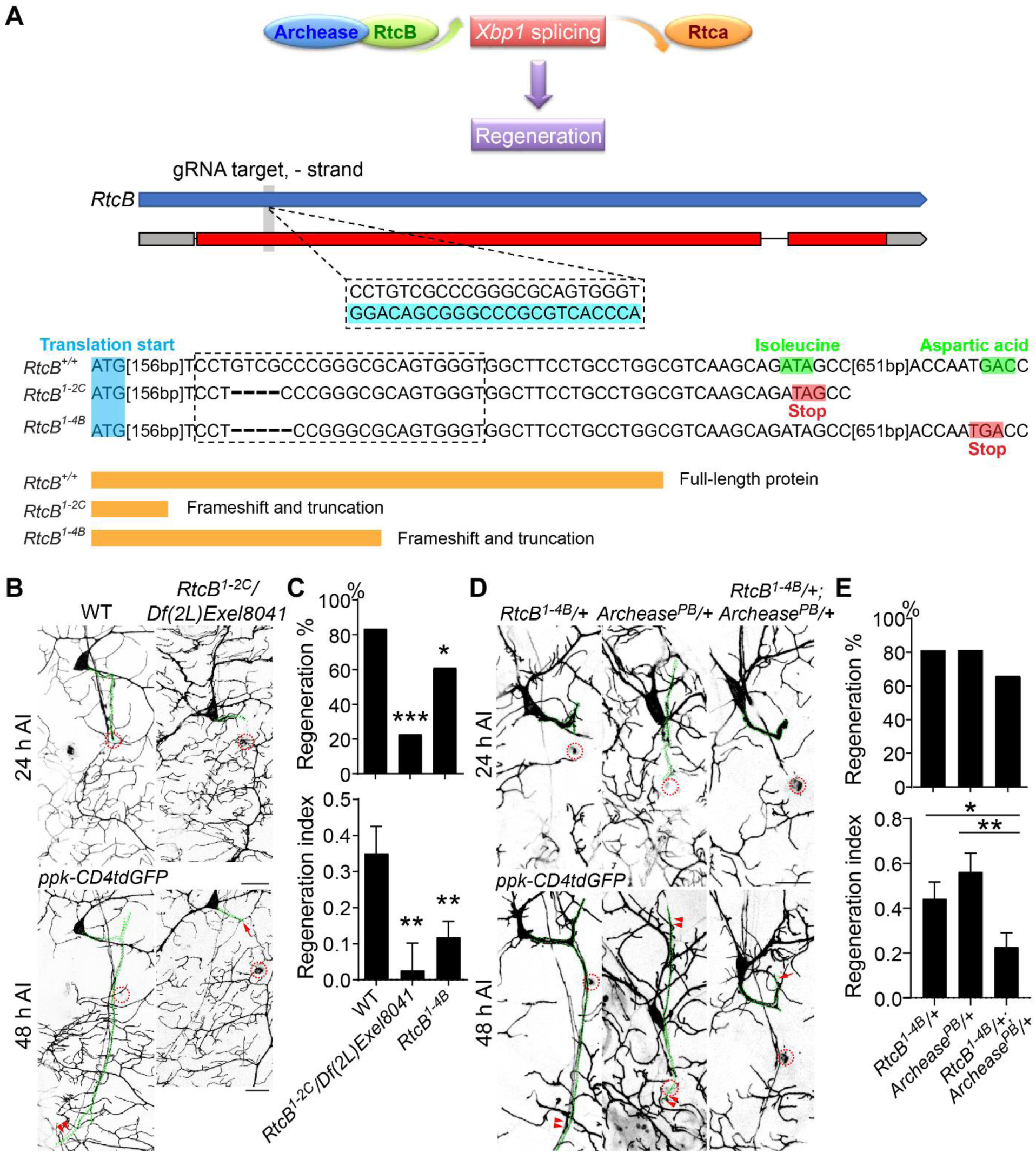
RtcB cooperates with its cofactor Archease to promote axon regeneration. (A) Generation of *RtcB*^1–^*^2C^* and *RtcB*^1–^*^4B^* mutant flies. Upper panel: proposed model for Rtca and RtcB regulating axon regeneration in opposite directions via competition to splice *Xbp1*. Middle panel: design of the gRNA targeting RtcB. Lower panel: CRISPR editing results in frameshift and truncates the protein. (B and C) *RtcB*^1–^*^2C^* and *RtcB*^1–^*^4B^* reduce axon regeneration. (B) C4da neuron axons were injured and regeneration was assessed at 48 h AI. Axons are highlighted by the dotted green line. The injury site is marked by the dashed circle, regenerating and non-regenerating axons are labeled by arrowheads and arrow, respectively. Scale bar, 20 μm. (C) Quantification of axon regeneration by regeneration percentage (upper panel, Fisher’s exact test, *p* < 0.0001, *p* = 0.0341) and regeneration index (lower panel, one-way ANOVA followed by Dunnett’s multiple comparisons test). *n =* 36, 22 and 54 neurons. (D and E) Compared with *RtcB*^1–^*^4B^* or *Archease^PB^* heterozygotes, axon regeneration is significantly decreased in *RtcB* and *Archease* transheterozygotes. (D) C4da axons were injured and regeneration was assessed at 48 h AI. Scale bar, 20 μm. (E) Quantification of axon regeneration percentage (upper panel, Fisher’s exact test) and regeneration index (lower panel, Kruskal-Wallis test followed by Dunn’s multiple comparisons test). *n =* 32, 27 and 47 neurons. \**p* < 0.05, \*\**p* < 0.01, \*\*\**p* < 0.001. See also S1 Figure.

### Inhibiting Rtca promotes axon regeneration and recovery of locomotor functions after SCI in adult mice

Our prior work showed that Rtca LoF not only enhances sensory neuron axon regeneration in the fly CNS, but also modestly promotes RGC axon regeneration after optic nerve crush in adult mice^8^. To further validate Rtca’s conserved role in impeding nerve regeneration in the mammalian CNS, we used the more clinically-relevant spinal cord injury (SCI) paradigm by performing dorsal over-hemisection at thoracic 7 (T7) vertebral level in adult mice (Figure 2A). Since C57BL/6 mice lack the ventral CST axons, our over-hemisection typically transects the majority of spinal cord area and lesions all CST axons^16^. We found that mice with a LoF allele of Rtca – *Rtca^Ins/Ins^* ^8^ exhibited significantly improved motor recovery as indicated by the increased Basso Mouse Scale (BMS) scores compared with WT mice over the 6 weeks of observation after SCI (Figures 2A and 2B). Surprisingly, the mutant mice showed signs of recovery as early as day 2 after injury, implying that Rtca regulates cell functions by diverse signaling pathways, such as neuroprotection and short-range sprouting. To further assess the locomotor recovery of the hindlimbs after SCI, we also performed the hindlimb grid walk and touch-grasping tests 4 and 6 weeks after SCI. *Rtca^Ins/Ins^* mice significantly surpassed WT in recovery by making fewer grid walk errors and having a higher grasping rate of hindlimbs (Figures 2C and 2D), suggesting that CST regeneration at least partly contributes to the functional recovery after SCI. Six weeks after SCI, we evaluated the regeneration of injured corticospinal tracts (CSTs) labeled by tracer biotin dextran amine (BDA). CSTs are important for controlling voluntary movements^17^, but particularly refractory to regeneration after axotomy^18–20^. Consistent with the behavioral data, *Rtca^Ins/Ins^*mice showed multiple regenerating CST axons extended beyond the injury site, with the longest axon reaching about 2-3 mm caudal to the lesion (Figures 2E-2G). In contrast, CST axons terminated at the lesion site in WT controls. We carefully checked the CST axons in the caudal spinal cord and confirmed that they meet the previously defined morphological criteria of regenerating axons^21^. Examination of reactive glial scar areas around the lesion site labeled by GFAP indicated that Rtca deletion did not alter glial scar size (S2 Figure). These data demonstrated that Rtca acts as a conserved anti-regeneration factor in the mammalian CNS.

**Figure 2.**
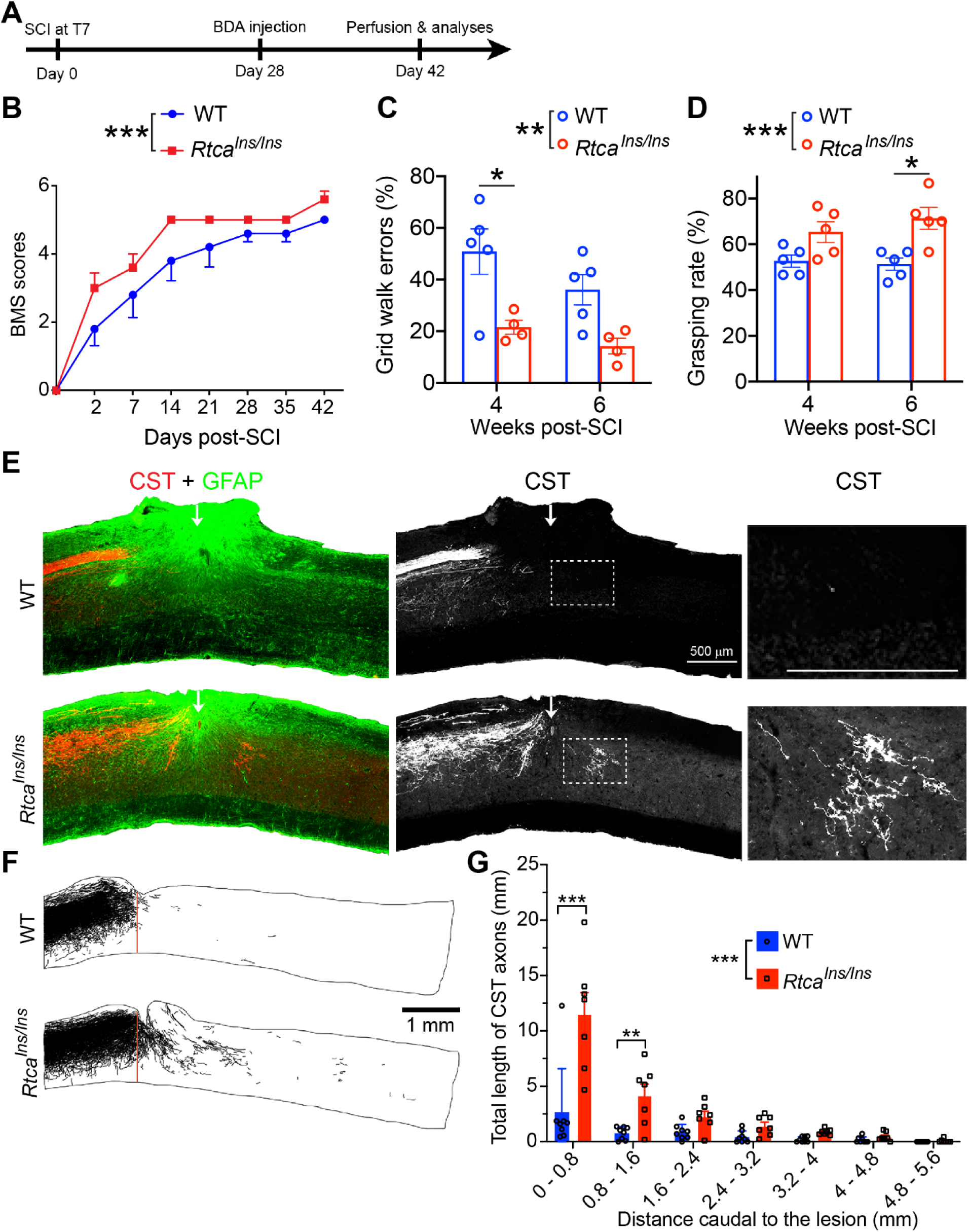
Rtca LoF enhances axon regeneration and recovery of locomotor functions after SCI in mice. (A) Schematic of the experimental protocol for mice SCI. BDA was injected to label CST axons at 28 days after over-hemisection at T7 and mice were perfused at 42 days after injury. (B) The BMS was conducted at multiple time points after SCI. Compared with WT, *Rtca^Ins/Ins^* mice showed better motor improvement and higher BMS scores over six weeks of observation. Two-way ANOVA. *n =* 5 and 5 mice. (C) Grid walk tests were performed at 4- and 6-weeks post SCI. *Rtca^Ins/Ins^* mice made fewer grid-walk errors than the WT mice, especially at 4 weeks after SCI. Two-way ANOVA followed by Sidak’s multiple comparisons test. *n =* 5 and 4 mice. (D) Grasping tests were performed at 4- and 6-weeks post SCI. *Rtca^Ins/Ins^* mice showed significantly increased hindlimb grasping rate, especially at 6 weeks after SCI. Two-way ANOVA followed by Sidak’s multiple comparisons test. *n =* 5 and 5 mice. (E) Representative images showing CST axons and GFAP^+^ lesion area. Scale bar, 500 μm. (F) Representative examples of camera lucida drawing of CST axons showing that Rtca LoF promotes axon regeneration after SCI. Scale bar, 1 mm. (G) Quantification of total CST axons regrowing beyond the injury site. There are more regrowing axons in *Rtca^Ins/Ins^* mice, especially within the 0-1.6 mm beyond the injury site. Two-way ANOVA followed by Bonferroni’s multiple comparisons test. *n =* 8 and 7 mice. \**p* < 0.05, \*\**p* < 0.01, \*\*\**p* < 0.001. See also S2 Figure.

### Rtca maintains proper expression and function of Piezo

Despite functioning as a potent regeneration inhibitor, how Rtca exerts its function is scarcely studied. To uncover potential Rtca effectors, we previously conducted transcriptome profiling of C4da neurons, comparing WT and *Rtca* LoF mutants – *Rtca^NP5057^* ^8^. By focusing on the genes upregulated in *Rtca* mutants, we identified a downstream pro-regeneration factor, ringer, which is suppressed by Rtca^9^. To uncover candidate regeneration inhibitors, we further queried the genes downregulated in *Rtca* mutants and found that *Piezo*’s transcript level is modestly reduced by 34% (Figure 3A). To test whether Piezo’s expression is affected by Rtca at the protein level, GFP was fused to the N-terminal of Piezo to generate GFP-Piezo fusion protein without disturbed Piezo’s channel function as previously described^11,22^. As a channel protein, Piezo was preferentially enriched on the plasma membrane in WT neurons. However, in *Rtca* mutant neurons, Piezo’s membranous enrichment was largely abolished (Figures 3B and 3C), as indicated by the decreased membrane/cytoplasm fluorescence intensity ratio (Figure 3D). To determine whether Rtca and Piezo act in the same genetic pathway, we performed transheterozygotes analysis for knockouts of *Piezo* (*PiezoKO*) and *Rtca* (*Rtca^Δ^*)^8^. Since both Rtca and Piezo suppress axon regeneration, we turned to C3da neurons which are incapable of regenerating their axons after axotomy. We found that while single heterozygotes of *Rtca^Δ^/+* or *PiezoKO/+* C3da neurons did not increase regeneration ability, *Rtca* and *Piezo* transheterozygotes (*Rtca^Δ^/+; PiezoKO/+*) possessed much stronger regeneration capacity, reaching the level seen in *Rtca* mutants or *PiezoKO* (Figures 3E and 3F), indicating that Rtca and Piezo do genetically interact to suppress axon regeneration and that Piezo likely lies downstream of Rtca.

**Figure 3.**
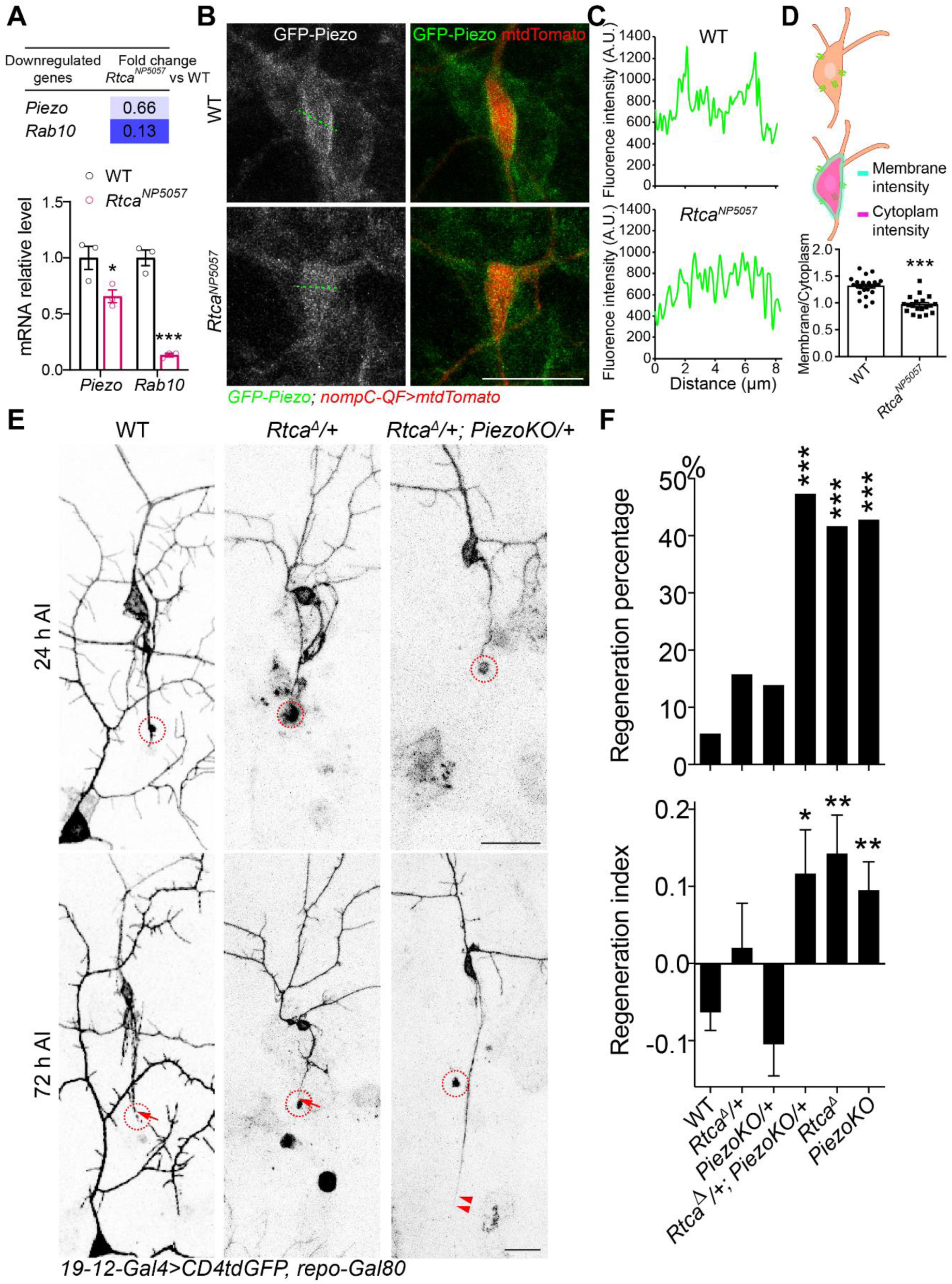
Rtca regulates Piezo’s expression and function. (A) RNA-seq analysis showing both Piezo and Rab10 are downregulated in *Rtca* mutant C4da neurons. The value of *Rtca* mutant is normalized to that of WT. Analyzed by unpaired *t*-test, *n =* 3, *p* = 0.0444, *p* = 2.68×10^−4^. (B) Representative images showing that Piezo is enriched on plasma membrane in WT but not in *Rtca* mutants. An example trace is shown in the dotted green line. Scale bar, 20 μm. (C) GFP-Piezo fluorescence intensity plot reveals that in WT neurons Piezo’s intensity is higher on the plasma membrane than in the cytoplasm. (D) Quantification of the membrane/cytoplasm ratio of GFP-Piezo fluorescence intensity. Unpaired *t*-test with Welch’s correction, *p* < 0.0001. *n =* 25 and 19 neurons. (E and F) Axon regeneration is significantly enhanced in *Rtca* and *Piezo* transheterozygotes. (E) C3da neuron axons were severed and their regeneration was assayed at 72 h AI. The injury site is outlined by the dashed circle, regenerating and non-regenerating axons are marked by arrowheads and arrow. Scale bar, 20 μm. (F) Quantification of axon regeneration shown in (E). Upper panel: regeneration percentage, data are analyzed by Fisher’s exact test, *p* = 0.3237, *p* = 0.2611, *p* = 0.0004, *p* = 0.0008, *p* < 0.0001. Lower panel: regeneration index, analyzed by one-way ANOVA followed by Dunnett’s multiple comparisons test. *n =* 37, 19, 36, 19, 24, and 49 neurons. \**p* < 0.05, \*\**p* < 0.01, \*\*\**p* < 0.001.

### Rab10 inhibits axon regeneration downstream of Rtca

Another gene revealed by RNA-seq is Rab10, a member of the diverse Rab family of small GTPases, whose transcription was decreased by 87% in *Rtca* mutant fly neurons (Figure 3A). To validate the decrease of Rab10 in *Rtca* mutants, we utilized a Rab10-EYFP knock-in fly stock^23^ to label the endogenous Rab10 and found Rab10 fluorescence intensity was significantly decreased in the cell bodies of injured C3da neurons in *Rtca^NP5057^* mutants (Figures 4A and 4B). Consistently, reduced Rab10 expression was observed after axotomy in cultured primary hippocampal neurons derived from *Rtca^Ins/Ins^* mice (Figures 4C and 4D). To determine if Rab10 acts to inhibit axon regeneration, we first knocked down Rab10 by RNAi in C3da neurons and found these neurons showed enhanced regeneration capacity (Figures 4E and 4F). Similarly, when we overexpressed Rab10^T23N^, a dominant-negative form of Rab10 (Rab10-DN) in C3da neurons, the regeneration percentage and regeneration index were also significantly increased (Figures 4E and 4F). Lastly, we knock out Rab10 and observed enhanced axon regrowth as well (Figures 4E and 4F). Moreover, we assessed axon regeneration in *Crag* LoF mutant (*Crag^GG^*^43^) neurons, as Crag operates as a Rab10-GEF responsible for Rab10’s activation^24^, and the regeneration capacity consistently increased (Figures 4E and 4F). The evidence collectively demonstrates that inactivating Rab10 promotes axon regrowth. On the other hand, we observed a decrease in regeneration capacity in C4da neurons expressing Rab10^Q68L^, a constitutively active form of Rab10 (Rab10-CA), compared to WT C4da neurons, which exhibited robust regeneration after axotomy (Figures 4G and 4H). Lastly, expressing Rab10-CA in *Rtca* mutant C3da neurons attenuated their enhanced regeneration ability (Figures 4E and 4F). Contrarily, while overexpressing Rtca in C4da neurons impaired regeneration, knocking down Rab10 in combination reversed the regeneration reduction (S3 Figure). Together, these data revealed that Rab10 functions downstream of Rtca to suppress axon regeneration.

**Figure 4.**
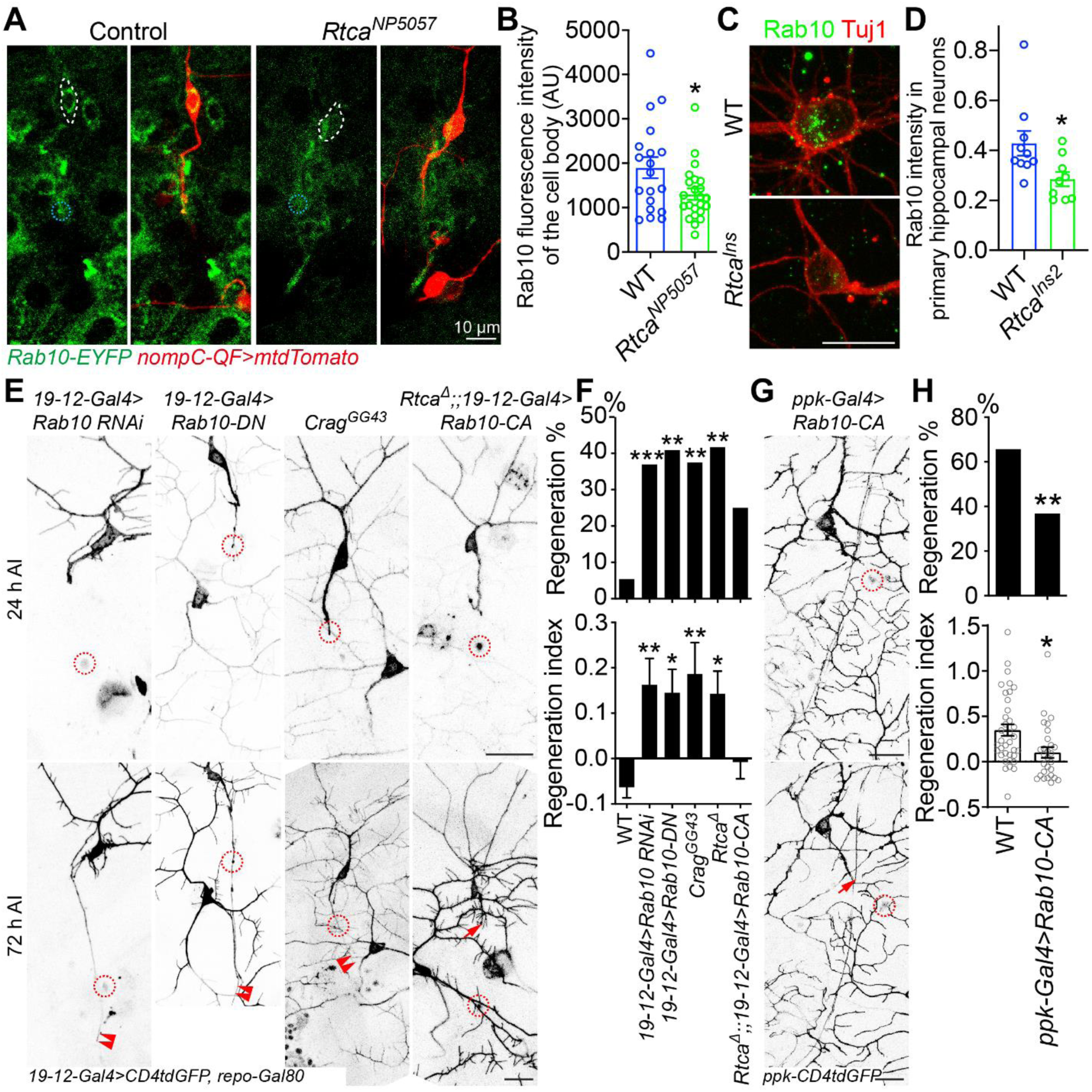
Rab10 operates downstream of Rtca to inhibit regeneration. (A) Rab10 fluorescence intensity is decreased in *Rtca* mutants. C3da cell bodies are marked by a dotted white line, and the injured site by a dotted blue circle. Scale bar, 10 μm. (B) Quantification of Rab10 fluorescence intensity shown in (A). *n =* 19 and 23 neurons. Analyzed by unpaired *t*-test, *p* = 0.0256. (C) Rab10 fluorescence intensity is reduced in injured primary hippocampal neurons cultured from *Rtca^Ins/Ins^* mice in microfluidic chambers. Scale bar, 20 μm. (D) Quantification of Rab10 intensity shown in (C). Rab 10 intensity is normalized to TUJ1. Analyzed by unpaired *t*-test, *p* = 0.0251. (E and F) Loss of functional Rab10 or inactivating Rab10 promotes axon regeneration, while overexpressing Rab10-CA in *Rtca* mutant attenuates the increased regeneration capacity. (E) C3da neuron axons were severed and their regeneration was assayed at 72 h AI. The injury site is marked by the dashed circle, regenerating and non-regenerating axons are marked by arrowheads and arrow. Scale bar, 20 μm. (F) Quantification of axon regeneration shown in (E). Upper panel: regeneration percentage, data are analyzed by Fisher’s exact test, *p* = 0.0006, *p* = 0.0013, *p* < 0.0001, *p* = 0.0043, *p* = 0.0008, *p* = 0.0837. Lower panel: regeneration index, analyzed by one-way ANOVA followed by Dunnett’s multiple comparisons test. *n =* 37, 46, 22, 33, 24, 24, 20 neurons. (G and H) Overexpressing Rab10-CA decreases regeneration capacity in regeneration-competent C4da neurons. (G) C4da neuron axons were injured and regeneration was assessed at 48 h AI. The injury site is marked by the dashed circle and non-regenerating axon is labeled by arrow. Scale bar, 20 μm. (H) Quantification of axon regeneration by regeneration percentage (Fisher’s exact test, *p* = 0.0267) and regeneration index (unpaired *t*-test with Welch’s correction, *p* = 0.0060). *n =* 37 and 28 neurons. \**p* < 0.05, \*\**p* < 0.01, \*\*\**p* < 0.001. See also S3 Figure.

### Rab10 mediates Piezo membrane targeting

Though Piezo is a well-documented regeneration suppressor, enriching at the injured axon tip to impede axon regrowth by triggering Ca^2+^ signaling^11^, how Piezo is recruited to and accumulated at the axon tip after injury remains unknown. Interestingly, Rab10 is involved in a wide range of activities such as vesicular transport in the endosomal/exocytosis recycling pathways^25^ and regulation of protein membrane insertion^26,27^. Since both Piezo and Rab10’s expression is regulated by Rtca, and they function in the same pathway as Rtca to inhibit regeneration, we asked if Piezo’s expression and localization are also affected by Rab10. We examined Piezo’s localization in Rab10 knockdown C3da neurons. In 45% of WT neurons, Piezo was enriched at the axon tip after injury. In comparison, only 30% of Rab10 knockdown neurons showed Piezo accumulation at the injured axon tip (Figures 5A and 5B). Consistently, the axon tip accumulation of phospho-CamKII (pCaMKII), which is activated by Piezo after axotomy and functions downstream of Piezo to impedes regeneration^11^, was also impaired in Rab10 knockdown neurons (S4 Figure). Since the Piezo intensity difference might be masked by the autofluorescence caused by injury, we looked at Piezo’s expression and localization in the soma and found Piezo’s membranous enrichment was impaired in Rab10 knockdown neurons after injury. In WT neurons, Piezo membrane intensity was significantly increased after injury, while its membrane/cytoplasm intensity ratio only showed a slight increase (Figures 5C-5E), implying Piezo can be properly localized to the membrane. Comparatively, there’s no such increase in Piezo’s membrane intensity in C3da neurons expressing Rab10 RNAi, and Piezo membrane/cytoplasm intensity ratio was significantly lower than WT at 24 h after injury (AI) (Figures 5C-5E), suggesting that Rab10 is critical for Piezo’s proper localization.

**Figure 5.**
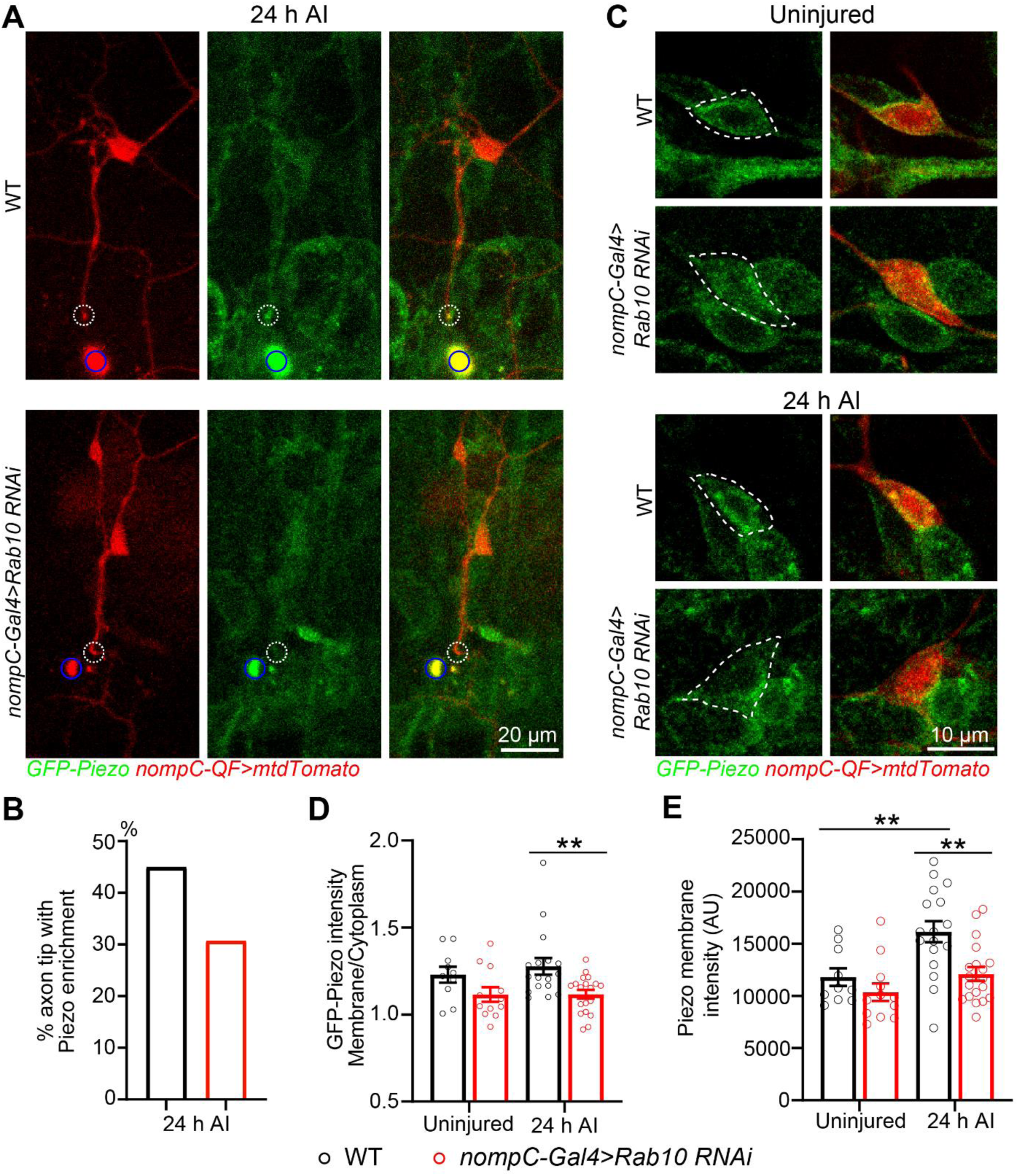
Rab10 mediates Piezo localization. (A) Representative images showing Piezo’s localization in injured neurons. The injury site is marked by blue circle and the growth cone is highlighted by the dotted white circle. Scale bar, 20 μm. (B) Quantification of the percentage of neurons with Piezo accumulating in the cut axonal tip. *n =* 40 and 26 neurons. Analyzed by Fisher’s exact test, *p* = 0.3072. (C-E) Loss of Rab10 reduces Piezo’s membrane localization after injury. (C) Piezo membrane intensity is increased after injury, while knocking down Rab10 impairs Piezo’s membrane enrichment. C3da neurons on one side of the larvae were injured and stained for GFP-Piezo and tdTomato (*nompC-QF>mtdTomato*), and the other side served as the uninjured control. (D) Quantification of Piezo membrane/cytoplasm ratio. (E) Quantification of Piezo membrane intensity. Uninjured: *n =* 10, 12 neurons; 24 h AI: *n =* 17, 19 neurons. Analyzed by two-way ANOVA followed by Sidak’s multiple comparisons test. \*\**p* < 0.01. See also S4 and S5 Figure.

A prior report showed that loss of Rab10 led to accumulated and swollen Rab5-positive early endosomes in *C. elegans* intestinal cells^12^. To examine whether Rab10 knockdown affects endosome trafficking in injured neurons in *Drosophila*, we expressed GFP-Rab5 in C3da neurons and assess GFP fluorescence intensity after axon injury. We noticed that compared with the dose control (*nompC-Gal4>BFP*), GFP-Rab5 intensity was significantly increased in Rab10 knockdown neurons at 24 h AI, while at 8 h and 48 h the intensity was comparable between the two groups (S5A and S5B Figure). Moreover, the GFP-Rab5 positive particle area was enlarged in Rab10 knockdown neurons (S5C Figure), suggesting Rab10 is implicated in endosome recycling after injury. Notably, Piezo is observed in GFP-Rab5 positive particles (S5D Figure).

### Rab10 is required for surface expression of integrin β1, which cooperates with Piezo to inhibit axon regeneration

We next explored if Rab10 regulates Piezo’s function via additional mechanisms. Prior reports showed that Piezo1’s activity is related to integrin-matrix interactions, and that integrin signaling pathways are essential for Piezo1’s activation upon membrane shear due to fluid flow in endothelial cells^28^. In turn, Piezo1 is important for the activity of Itgb1 in epithelial cells^29^. Moreover, it was previously reported that Rab11 induces Itgb1 surface expression^30^. Considering the synergy and partial redundancy among Rabs in transporting and trafficking^24^, we reasoned that Rab10 may play a role in regulating the localization of mys, the fly ortholog of Itgb1, in response to neural injury. We interrogated the surface expression of mys in C3da neurons and detected an over 1.5-fold increase of surface mys fluorescence intensity in WT neurons at 24 h AI compared to the uninjured control (Figures 6A and 5B). However, mys surface expression remained unchanged after injury in neurons overexpressing Rab10-DN (Figures 6A and 5B), indicating that the surface expression of mys is increased in response to axotomy in a Rab10-dependent manner. We then performed immunostaining and found that in WT neurons, Piezo partially colocalized with mys on the membrane and their colocalization modestly increased after axon injury (S6A Figure), coinciding with the increase of Piezo membrane intensity. Interestingly, in C3da neurons expressing mys RNAi, Piezo’s membrane/cytoplasm intensity ratio was decreased before and after injury (Figures 6C and 6D), suggesting that mys is required for regulating Piezo’s anchorage and function. Meanwhile, the percentage of axon tips showing Piezo enrichment at 24 h AI was substantially reduced in mys knockdown neurons compared with WT (S6B Figure).

**Figure 6.**
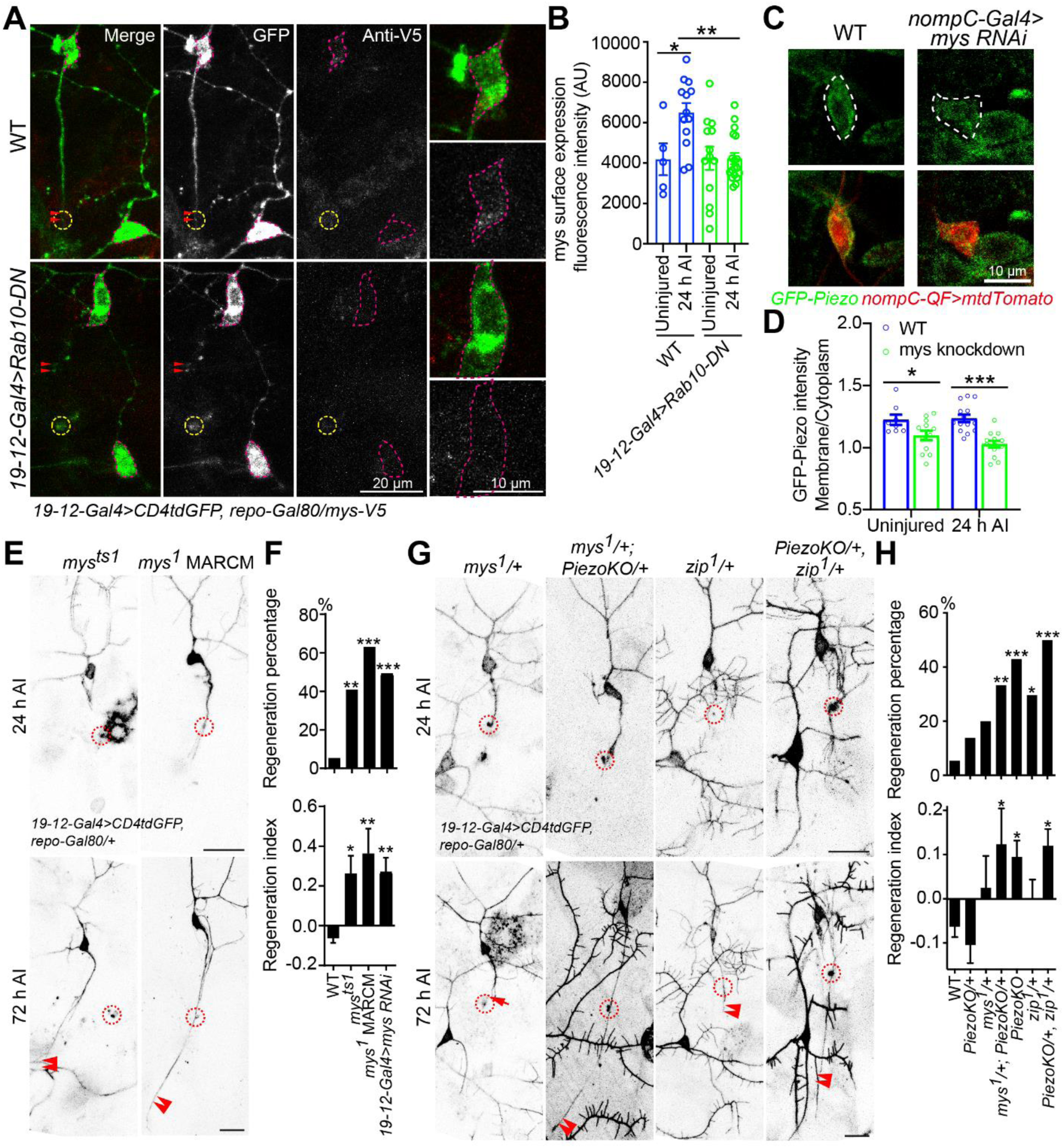
mys genetically interacts with Piezo to suppress axon regrowth. (A) mys surface expression is increased in WT neurons after injury but not in neurons expressing Rab10-DN. C3da cell bodies are marked by magenta dotted lines, and the injury sites are marked by dotted yellow circles. Axon tips are marked by double arrowheads. Scale bar, 20 μm (left panel) and 10 μm (right panel, enlarged view). (B) Quantification of mys surface expression before and after injury. Analyzed by two-way ANOVA followed by Holm-Sidak’s multiple comparisons test. *n =* 5, 13, 13, and 18 neurons. (C) Knocking down *mys* reduces Piezo’s membrane/cytoplasm intensity ratio in C3da neurons. C3da cell bodies are marked by a dotted white line. Scale bar, 10 μm. (D) Quantification of GFP-Piezo fluorescence. Uninjured: *n =* 8, 12 neurons; 24 h AI: *n =* 14 and 13 neurons. Analyzed by two-way ANOVA followed by Holm-Sidak’s multiple comparisons test. (E and F) Axon regeneration is enhanced in *mys* mutants and mys knockdown. C3da neuron axons were severed and their regeneration was assayed at 72 h AI (E). The injury site is marked by the dashed circle and regenerating axon is labeled by arrowheads. Scale bar, 20 μm. Quantification of axon regeneration percentage and index shown in (F). Upper panel: regeneration percentage, data are analyzed by Fisher’s exact test, *p* = 0.0013, *p* < 0.0001, *p* < 0.0001. Lower panel: regeneration index, analyzed by one-way ANOVA followed by Dunnett’s multiple comparisons test. *n =* 37, 22, 16 and 47 neurons. (G and H) Axon regeneration is significantly enhanced in *mys* and *Piezo* transheterozygotes, as well as in *zip* and *Piezo* transheterozygotes. C3da neuron axons were severed and their regeneration was assayed at 72 h AI (G). The injury site is marked by the dashed circle, regenerating and non-regenerating axons are marked by arrowheads and arrow. Scale bar, 20 μm. Quantification of axon regeneration shown in (H). Upper panel: regeneration percentage, data are analyzed by Fisher’s exact test, *p* = 0.2611, *p* = 0.1372, *p* = 0.0044, *p* < 0.0001, *p* = 0.0133, *p* < 0.0001. Lower panel: regeneration index, analyzed by one-way ANOVA followed by Dunnett’s multiple comparisons test. *n =* 37, 36, 15, 33, 49, 27, and 40 neurons. \**p* < 0.05, \*\**p* < 0.01, \*\*\**p* < 0.001. See also S6 and S7 Figure.

Piezo’s channel activity is indispensable for its role as a regeneration suppressor^11^, therefore, we reasoned that mys, which is important for Piezo’s membrane targeting, is involved in the anti-regeneration pathway. We first examined axon regeneration in mys knockdown and two *mys* mutants, *mys^ts^*^1^ (a temperature-sensitive hypomorphic allele^31^) and *mys*^1^ (an amorphic allele^31^). Increased C3da neuron axon regrowth was observed in *mys^ts^*^1^ mutants and *mys*^1^ MARCM clones as well as mys knockdown (Figures 6E and 6F), confirming that LoF of mys promotes drastic axon regeneration. Then we expressed mys along with Rab10 RNAi in C3da neurons, and found it substantially attenuated the regeneration capacity in Rab10 knockdown neurons, suggesting that mys acts downstream of Rab10 to suppress axon regeneration (S7 Figure). To determine if Piezo and mys function in the same genetic pathway, we generated transheterozygotes, *mys*^1^*/+; PiezoKO/+*, and found robust axon regeneration after injury (Figures 6G and 6H). This suggests that mys functions in the same genetic pathway as Piezo to suppress axon regeneration. The transheterozygotes of *Piezo* and *zip*, the fly ortholog of non-muscle myosin II (NM2, an important component in forming cell-matrix adhesions and suppressing regeneration^32^) also displayed stronger regeneration capacity (Figures 6G and 6H), demonstrating that the integrin-matrix interaction is vital for Piezo’s anti-regeneration function.

### Rab10 and Itgb1 inhibit axon regeneration in the adult mammalian CNS

Our findings reported above uncovered an anti-regeneration pathway downstream of Rtca. To determine if Rab10 and Itgb1/mys, two critical genes in this pathway, also inhibit regeneration in mammals, we injected AAV2 expressing Rab10-DN into the retina of adult mice and crushed the optic nerves 7 days after AAV2 application (Figure 7A). At 21 days after injury, we observed significantly enhanced axon regeneration in mice treated with AAV2-Rab10-DN but not in the controls treated with AAV2-GFP (Figures 7B and 7C). This result proves that Rab10 does impede CNS axon regeneration in adult rodents. Furthermore, we obtained the *Itgb1^f/f^* mice^33^ and deleted *Itgb1* gene selectively in retinal cells, including RGCs, by intravitreal injection of AAV2 expressing Cre. We found that loss of Itgb1 promoted drastic axon regeneration of axotomized RGCs 21 days after optic nerve crush (Figures 7F and G). Interestingly, ablating Itgb1 modestly protected RGCs from injury-induced cell death, which is reminiscent of the phenotype observed in neurons expressing Rab10-DN (Figures 7D, 7E, 7H and 7I). Therefore, deleting either Rab10 or Itgb1 stimulates significant axon regeneration of injured CNS neurons in adult mammals.

**Figure 7.**
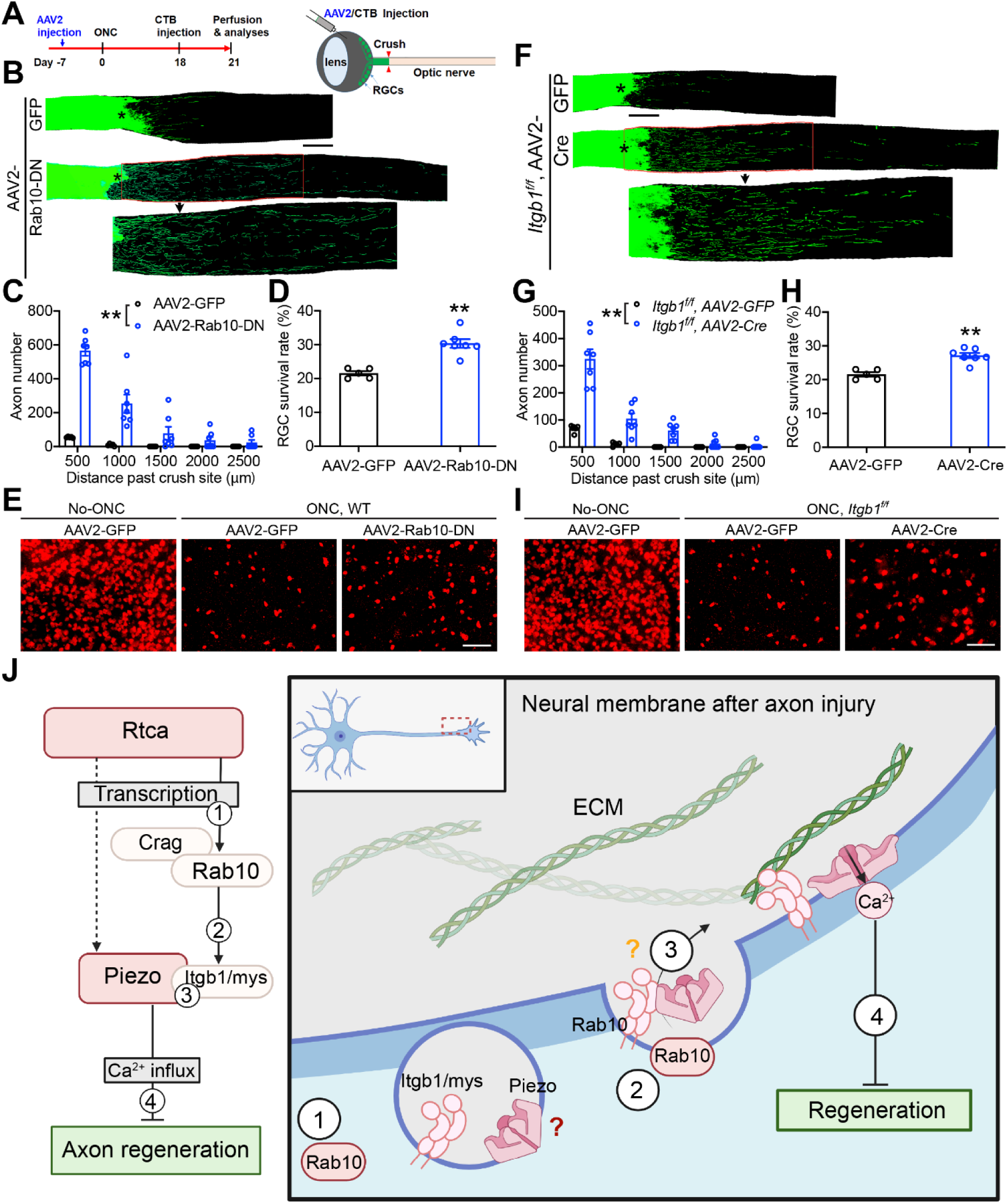
Rab10 and Itgb1 impede CNS nerve regeneration in adult mice. (A) Schematic of experimental protocols for the mice with optic nerve crush. AAV2-GFP, AAV2-Rab10-DN or AAV2-Cre was injected into the retina intravitreally 7 days before injury. CTB was injected at 18 days after optic nerve crush and mice were perfused at 21 days after injury. (B-E) Loss of functional Rab10 promotes axon regeneration after optic nerve crush. (B) Representative images of optic nerve after crush in control and Rab10-DN overexpression. Scale bar, 100 μm. (C) Quantification of the axon numbers extending beyond the injury cite. (D) Quantification of RGC survival rate after injury. (E) Representative images of the RGCs. *n =* 5 and 7 mice. Scale bar, 50 μm. Data are analyzed by two-way ANOVA (C) or unpaired *t*-test (D). (F-I) Itgb1 conditional knockout enhances RGC axon regrowth in the mice CNS. (F) Representative images of optic nerve after crush in control and Itgb1 cKO mice. Scale bar, 100 μm. (G) Quantification of the axon numbers extending beyond the injury cite. (H) Quantification of RGC survival rate after injury. (I) Representative images of the RGCs. *n =* 4 and 7 mice. Scale bar, 50 μm. Data are analyzed by two-way ANOVA (G) or unpaired *t*-test (H). (J) Schematic drawing depicting how Rtca regulates axon regeneration by mediating Piezo’s localization. It remains to be determined that if Piezo and Itgb1/mys colocalize in Rab10-mediated versicles (red question mark) and if Itgb1/mys physically interact with Piezo to recruit Piezo to the membrane (orange question mark). \*\**p* < 0.01. See also S8 Figure.

To conclude, we have demonstrated that after injury, Rab10 acts downstream of Rtca to upregulate the surface expression of Itgb1/mys, which is critical for Piezo’s proper localization to the membrane and inhibition of axon regeneration. In the absence of Rtca, Rab10 is dramatically downregulated. As a result, the axotomy-induced upregulation of surface Itgb1/mys is abolished, decreasing membranous Piezo and perturbing its anchorage and activation, thus facilitating axon regeneration (Figure 7J).

## Discussion

Piezo has been documented as a mechanosensor, able to detect the rigidity and topography of the extracellular matrix in response to the environmental stiffness after axon injury and during regeneration^10^. Prior work has demonstrated that Piezo mediates Ca^2+^ influx and triggers downstream pathways in response to injury. Different from L-type Ca^2+^ channels which mediated global calcium transients and promote regeneration^34^, the Ca^2+^ influx via mechanosensitive channel exerts an inhibitory effect locally at the axon tips, preventing axon from regrowth^11,35^. But how Piezo is recruited to the injured axon tip remained to be elucidated. This work revealed an evolutionarily conserved inhibitory pathway for regeneration by regulating Piezo’s localization and function. After axotomy, Rtca controls the stress-induced mRNA splicing. Rab10 functions downstream of Rtca to increase the surface expression of mys and mediates Piezo’s enrichment and anchorage on the membrane. Piezo then senses the mechanical force in the microenvironment, converting it into intracellular signals that prevent axons from regenerating (Figure 7J).

Our findings link the injury-induced RNA repair/splicing to membrane trafficking. During cellular stress, Rtca is recruited to mediate RNA metabolism. Prior results demonstrated that Rtca is an anti-regeneration factor^8^. Here we further demonstrated that in flies, Rtca and RtcB function oppositely in axon regeneration. Whereas RtcB cooperates with Archease to unconventionally splice *Xbp1*, generating an activated *Xbp1*^14^, which benefits myelin removal and axon regeneration^36^, Rtca antagonizes the reaction to slow down *Xbp1* splicing, impeding axon regrowth^8^. However, many of the downstream effectors in the Rtca pathway during regeneration have not been explored, dependent or independent of Xbp1 splicing. *Rtca* mutant RNA-seq data showed that in sensory neurons, the expression levels of more than 200 genes were altered compared to WT. Our prior work showed that the microtubule-associated protein (MAP) ringer/TPPP3 is highly upregulated in *Rtca* mutants, which interacts with futsch/MAP1B and HDAC6 to modulate microtubule dynamics and promotes axon regeneration (S8 Figure)^9^. Interestingly, Gene Ontology (GO) analysis reflected that the pathways regulating extracellular matrix and plasma membrane topped the list of those affected in *Rtca* mutants^9^. Intriguingly, Rab10, a critical regulator of membrane organization and composition by controlling post-Golgi vesicle trafficking and endosomal sorting^25^, is among the genes downregulated most strikingly in the absence of Rtca.

Rab10 has been reported to promote axon and dendrite outgrowth during development via mediating membrane trafficking^26,27,37^, which is essential for localizing trans-membrane proteins (*e.g.*, DMA-1 and HPO-30) to the membrane^37,38^. Upon neural injury, several proteins related to endosome sorting and membrane trafficking are upregulated, including Rab4 and rabaptin-5^39^, suggesting that this pathway dynamically regulates injury-induced cellular stress. Here we showed that after axon injury, Rab10 depletion interferes early endosome recycling, leads to delayed endosome transporting and accumulated Rab5^+^ endosomes, which may impair the biological processes depending on the endosome network and in responding to axotomy. One such component, integrin, whose expression and distribution are mainly controlled by endocytosis and recycling^40^, is severely affected in Rab10 knockdown neurons. Previously, Rab10 was shown to directly interact with Itgb1 and colocalize with integrins in endosomal versicles^41^. We found that in intact neurons, the surface expression of mys is comparable between WT and Rab10-DN, while after injury mys surface expression increases in WT but not Rab10-DN neurons. This suggests that Rab10 is necessary for mys’ quick response to injury-induced stress, although other Rab members may compensate for Rab10 and properly localize mys to the membrane.

Integrins link the interactions between cells and the surrounding matrix, and are potentially critical for wound healing^42^ and axonal growth/guidance^43^. Nevertheless, the role of integrins during axon outgrowth and regrowth is controversial. Although integrins α7 and α9 act as pro-regeneration factors in rodents^44,45^, Itgb1 suppresses axon growth in the spinal cord^46^. Moreover, Kingston *et. al.* found that loss of integrin β3 not only protects RGCs from cell death, but also promotes RGC axon regeneration after optic nerve injury^47^. In comparison, our findings showed that deleting mys/Itgb1 drastically enhances axon regeneration in both flies and mice while increasing RGC survival in mice moderately. Given that α and β integrin subunits interact with each other and can form functional heterodimers^48^, cells may dynamically adjust the membrane expression of different integrin subunits corresponding to the complex environment to mediate regeneration after injury. However, further studies are needed to determine the precise compositions of integrin subunits for promoting cell survival, axon regeneration, and functional recovery.

Mechanosensation is one of the main functions of integrin-mediated cells and extracellular matrix adhesion, and integrin functions as a mechanosensor itself^49^. Piezo is associated with integrin-mediated mechanotransduction^48^. Prior report has shown that upon force, Piezo1’s recruitment and stabilization in adhesions correlates with the enrichment of β1 integrins while anticorrelates with β3 integrins, suggesting that integrin may bind in a complex with Piezo to regulate Piezo’s recruitment^28^. Here we showed that mys interacts with Piezo to regulate Piezo’s localization and function after injury. Considering Piezo is observed in Rab5-labeled early endosomes, increasing Piezo’s membrane concentration at the cut axonal tip through the endocytic recycling pathway together with integrins is potentially an energy-saving strategy adopted by injured neurons. Notably, Piezo’s expression and localization are altered in the absence of Rtca even before injury, so we are unable to rule out the possibility that Rtca directly regulates Piezo expression. Interestingly, zip/NM2 also functions in the same genetic pathway as Piezo to inhibit axon regeneration, suggesting that cell and extracellular matrix adhesion is necessary for Piezo’s function during regrowth. Our lab previously found that Piezo operates to convert environment stiffness/rigidity into intracellular signaling and activates the Atr-Chek1 pathway to regulate axon regeneration^10^. These collectively suggested that after injury, Rtca-Rab10-mys pathway facilitates Piezo’s enchainment on the axon tip, where Piezo senses the stiffness of the matrix and triggers anti-regeneration signaling. The fact that the machinery is conserved from *Drosophila* to mice is encouraging, enabling the screening with the fly model for the antagonists of the pathways to promote nerve repair.

## Acknowledgement

We thank Bloomington Stock Center and VDRC for fly stocks; Feng Zhang for helping with the RNAseq data analysis, members of the Song lab for helpful discussions. This work was supported by NIH grants (1R01NS107392; 1R01NS126541) and Pennsylvania Department of Health grant (4100088540) to Y.S., and by research grants to S.L. from NIH (R01NS105961, 1R01NS122813, and 1R01EY033652), Shriners Research Foundation (85133-PHI-21), and Wings For Life Spinal Cord Research Foundation (WFL-US-09/21-249).

## Author contributions

Experimental design, Q.W., L.M., S.L. and Y.S.; Methodology, Q.W., L.M., S.L. and Y.S.; Data collection and analysis, Q.W., L.M., S.W., H.N.N., E.J.M.V., J.P., S.J.O., S.L. and Y.S.; Writing – Original Draft, Q.W., L.M. and Y.S.; Writing – Review & Editing, J.P. and S.L.; Funding Acquisition, L.S. and Y.S.; Supervision, L.S. and Y.S.

## Author information

The authors declare no competing interests. Correspondence and requests for materials should be addressed to Y.S.

## Data availability

All data supporting the findings of this study are provided within the paper and its Supplementary Information. Source data are provided with this paper.

## Methods

### Fly stocks

*19-12-Gal4*^50^*, repo-Gal80*^51^*, ppk-CD4-tdGFP*^52^*, ppk-Gal4*^52^*, UAS-CD4tdGFP*^52^*, nompC-Gal4*^53^, *nompC-QF*^53^, *QUAS-mtdTomato*^54^ , *QUAS-mCD8GFP*^54^, GFP-Piezo^11^, *Rtca^NP5057^* ^8^, *Rtca^Δ^* ^8^ *,UAS-Rtca*^8^ and *PiezoKO*^11^, *Rab10^EYFP^* ^23^, *UAS-YFP.Rab10.Q68L* (Rab10-CA)^55^, *UAS-YFP.Rab10.T23N* (Rab10-DN)^55^, *mys*^1 31^, *mys^ts^*^1 31^ and *zip*^1 56^ have been previously described. *UAS-Rab10 RNAi* (BL#26289), *UAS-GFP-Rab5* (BL#43336)*, UAS-mys RNAi* (BL#33642), *UAS-mys* (BL#68158), *UAS-mRFP* (BL#7119) and *UAS-BFP* (BL#56807) were from Bloomington stock center and *mys^fTRG00932.sfGFP-TVPTBF^* (*mys-V5*) (v318285) was from VDRC. Piezo-Flag is a generous gift from Xiang Yang. To generate the *RtcB*^1–^*^2C^* and *RtcB*^1–^*^4B^* alleles, gRNA was cloned into the pU6-3-gRNA vector^57^ and injected into Cas9 expressing flies (Rainbow Transgenic Flies, Inc). Randomly selected male and female larvae were used.

### Mice

All studies and procedures involving animal subjects were performed under the approval of the Institutional Animal Care and Use Committee (IACUC) at Temple University. C57BL/6J and *Itgb1^f/f^* mice were obtained from the Jackson Laboratory. *Rtca^Ins/Ins^* mice was previously described^8^. Adult mice (10 weeks old) with the same sex were randomly assigned to experimental groups. All mice were housed in an animal facility and maintained in a temperature and light controlled environment with an alternating 12-hour light/dark cycle. The animals had no prior history of drug administration, surgery or behavioral testing.

### Live imaging in flies

Live imaging was performed as described^58^. At the appropriate time, a single larva was anesthetized with ether and mounted on slide with 90% glycerol under coverslips sealed with grease. The larva was then imaged with a Zeiss LSM 880 microscope and returned to grape juice agar plates after imaging sessions.

### Sensory axon lesion in *Drosophila*

Da sensory neuron injuring and imaging was performed in live fly larvae as described^7,8,59^. Embryos were collected for 2-24 hours on yeasted grape juice agar plates and were incubated at 25 °C. At 48 or 72 h after egg laying, larvae were anesthetized and mounted on glass slide. Axon was severed with a focused 930-nm two-photon laser and the lesion was confirmed 24 h AI. At 48 h AI (for C4da) or 72 h AI (for C3da), axon regeneration was assayed with “regeneration percentage” to quantify the percent of regenerating axons and “regeneration index” to normalize the axon elongation to larvae growth (“axon length”/“distance between the cell body and the axon converging point”), according to published methods^7,8^. An axon was defined as regenerating only when it obviously regenerated beyond the retracted axon stem, and this was independently assessed of regeneration index. The regeneration parameters from various genotypes were compared with that of the WT if not noted otherwise, and only those with significant differences were labeled with the asterisks.

### Fly immunohistochemistry

Third instar larvae were dissected and their body walls were fixed with 4% PFA according to standard protocols. The tissue was blocked in blocking buffer (PBS + 0.3% Triton X-100 + 5% normal donkey serum) and then incubated with primary antibodies. For mys surface staining, PBS + 5% normal donkey serum was used for blocking the samples and diluting antibodies. The following primary antibodies were used: rabbit anti-GFP (1:1000, Abcam, ab290), chicken anti-GFP (1:1000, Abcam, ab13970), rabbit anti-RFP (1:1000, Rockland Immunochemicals, 600-401-379), mouse anti-mys (1:1000, DSHB, CF.6G11), mouse anti-V5 (1:400, Thermofisher, MA5-15253) and anti-phospho-CamKII alpha/bata/delta (Thr305) (1:400, Thermofisher, PA5-37832). Fluorescence-conjugated secondary antibodies (1:200, Jackson ImmunoResearch) were used for immunohistochemistry. Larval were mounted in VECTASHIELD Antifade Mounting Medium. To assess Piezo membrane/cytoplasm ratio, the boundary of the C3da neuron is defined by membrane-targeting tdTomato (myristoylated and palmitoylated Tomato^54^, expressed under the control of *nompC-QF*), then the membranous and cytoplasmic area of the neuron were outlined as described in Figure 3D and GFP fluorescence was measured by ImageJ.

### SCI and axon evaluation in adult mice

All studies and procedures involving animal subjects were performed under the approval of the Institutional Animal Care and Use Committee (IACUC) at Temple University. To lesion the spinal cord of WT (controls) and *Rtca^Ins/Ins^* mice (10 weeks old, C57BL/6 background), we exposed the dorsal spinal cord by T6-7 laminectomy. A dorsal over-hemisection (1 mm in depth, and approximately 1.5 mm in dorsoventral diameter) was performed at T7 with a 30-gauge needle and microscissors to completely sever the dorsal spinal cord, including all the CST axons. The lesion depth of 1 mm was ensured by passing a marked 30-gauge needle at least 5 times across the dorsal spinal cord. Four weeks after SCI, the mice received BDA (10 kDa) tracer injections into 5 sites of the sensorimotor cortex (anterior-posterior coordinates from Bregma in mm: 1.0, 0.5, 0, -0.5, - 1.0, all at 1.0 mm lateral and at a depth of 1.0 mm). Mice were perfused 2 weeks after BDA injection and fixed spinal cords were dissected for histology.

To compare axon numbers in the caudal spinal cord between WT and *Rtca^Ins/Ins^* groups, we determined the length of BDA-labeled CST axons in all parasagittal sections of the spinal cord from 0 to 4 mm caudal to the lesion epicenter in each animal. The injury center was determined as the midpoint of histological abnormalities produced by lesion cavitation, reactive astrocytes, and morphological changes of injured axons. The CST axons caudal to the lesion were traced manually in each of the parasagittal sections and their total length inside of several bin boxes at 0.8, 1.6, 2.4, 3.2, and 4.0 mm caudal to lesion center was measured with Photoshop and Image J software. Tissue sections were immunostained with rat anti-GFAP antibody (1:50, ThermoFisher, 13-0300). We measured the GFAP^+^ dense scar tissue areas and the GFAP^+^ reactive astrocyte areas from multiple parasagittal sections of the lesioned spinal cord in each animal. We defined the former as the areas of densely overlapped GFAP^+^ astrocytic processes around the lesion epicenter and the latter as an obvious increase of GFAP immunoreactivity surrounding the scar tissues.

### Behavioral tests in mice with SCI

To determine functional recovery in *Rtca^Ins/Ins^* mice, we evaluated locomotion alterations during 6 weeks of survival by measuring multiple behavioral tests. The BMS scores were evaluated while the mouse was walking in an open field and confirmed from digital video records. The grid walk errors were counted from videotapes played at a slow speed (4 separate trials per test) and averaged from different trials. The contact-evoked grasping rate was measured by lowering the hindpaws toward a wire cage lid and determining the percent of times that it was grasped successfully. These behavioral tests were performed blindly by two persons who were unaware of animal identifications.

### Optic nerve crush injury, tracer injection, and evaluation of axon regeneration and RGC survival

To access the optic nerve, we used microscissors and dull/angled forceps to gently push away tissues near the eye and avoided damaging the orbital venous sinus. Fine angled forceps were employed to crush the optic nerve for 10 seconds with a consistent pressure at ∼1 mm behind the optic disc. To preserve the retinal blood supply, we were careful not to damage the underlying ophthalmic artery. To upregulate Rab10-DN in retina, we injected AAV2-GFP or AAV2-Rab10-DN (2 × 10^12^ genomic copy/mL) intravitreally in adult WT mice and crushed the optic nerves 7 days after AAV2 injection. To delete Itgb1 in the retina of adult *Itgb1^f/f^* mice, 7 days before optic nerve crush, we injected AAV2-GFP or AAV2-Cre (2 × 10^12^ genomic copy/ml) intravitreally at the crush side. At 18 days after injury, we labeled regenerating axons by anterogradely injecting Alexa488-conjugated CTB tracer (2 μL per mouse, 2.5 μg/μL) into vitreous with a micropipette. Three days after the CTB injection, mice were perfused with 4% PFA and optic nerves and retinas were collected. Fixed optic nerves containing injury sites were sectioned longitudinally (10 μm) and 5 representative sections were selected from each optic nerve by visualizing the most regenerating axons along the sections. We counted regenerating axons crossing several lines past the lesion and calculated their number by dividing the axon number by the nerve size at each distance. Following 2-hour post-fixation in the same PFA and overnight incubation in 30% sucrose, we immunostained the whole-mounted retina with an antibody for Tuj1 and examined the number of surviving RGCs in retina ipsilateral to injury. After quantifying the average number of Tuj1^+^ cells per field, we obtained the total number of viable RGCs by multiplying the figure by retinal area, as reported previously^16^.

### Neuron culture and immunostaining

Primary neuron culture was performed as previously described^11^. In brief, we dissociated primary mouse hippocampal neurons from postnatal day 0 (P0) mice and seeded the cells into our microfluidics devices. The neurons were cultured for 7 days (DIV 7) and the axons were injured via vacuum aspiration. The neurons were fixed and immunoassayed against class III beta-tubulin – mouse anti-Tuj1 (1:500, BioLegend, 801202) and Rab10 – rabbit anti-Rab10 (1:100, Cell Signaling Technology, 8127). The fixed samples were imaged with a Zeiss Celldiscoverer 7 confocal microscope and Rab10’s fluorescence intensity was quantified.

### RNA-seq data analysis

Analysis of RNA-seq data was performed as described previously^60^. Alignments of the sequencing data of three WT and three knockout samples were performed using STAR v2.5.2a to Ensembl *Drosophila* reference genome BDGP6.46^61^. The multimapping and chimeric alignments were discarded, and only uniquely mapped reads were quantified at the gene level and summarized to gene counts using STAR-quantMode (GeneCounts). Differential gene expression analysis between WT and mutant samples was performed using DESeq2 (v1.36.0) in R (v3.6.0) after genes whose average counts were lower than the 10 were discarded ^62^. Normalized gene counts of Rab10 and Piezo calculated by DESeq2 were used for visualization.

### Statistical Analysis

The sample sizes are similar to those reported in previous publications and the statistical analyses were done afterward without interim data analysis. The values of “*n*” (sample size) are provided in the figure legends. All data were collected and processed randomly. Data are expressed as mean ± SEM in bar graphs, if not mentioned otherwise. No data points were excluded. Two-tailed unpaired Student’s *t*-test was performed for comparison between two groups of samples. One-way ANOVA followed by multiple comparison test was performed for comparisons among three or more groups of samples. Two-way ANOVA followed by multiple comparison test was performed for comparisons between two or more curves. Fisher’s exact test was used to compare the percentage, each group was compared with the first row of dataset. All the mutants and RNAis were compared to WT unless specified otherwise. Statistical significance was assigned, \**p* < 0.05, \*\**p* < 0.01, \*\*\**p* < 0.001.

## Notes

### Competing Interest Statement

The authors have declared no competing interest.

