## Supplemental figures for "Targeting and anchoring the mechanosensitive ion channel Piezo to facilitate its inhibition of axon regeneration"

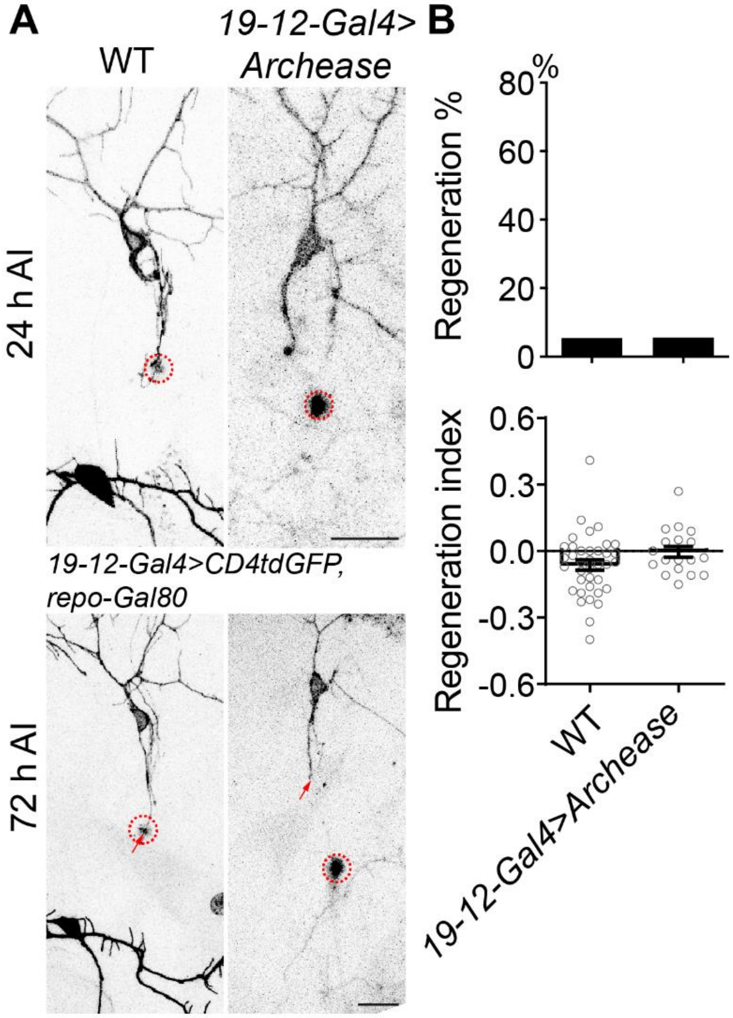


**S1 Figure. Overexpressing RtcB but not Archease is sufficient to increase regeneration, related to Figure 1.**

(A) C3da neuron axons were injured and assayed at 72 h AI. The injury site is marked by the dashed circle, regenerating axon is marked by arrowheads and non-regenerating axon is labeled by arrow.

(B) Quantification of axon regeneration by regeneration percentage (upper panel, Fisher's exact test, $p > 0.9999$, $p = 0.0123$) and regeneration index (lower panel, one-way ANOVA followed by Dunnett's multiple comparisons test). $n = 37$, 18 and 36 neurons. Scale bar, 20 μm.

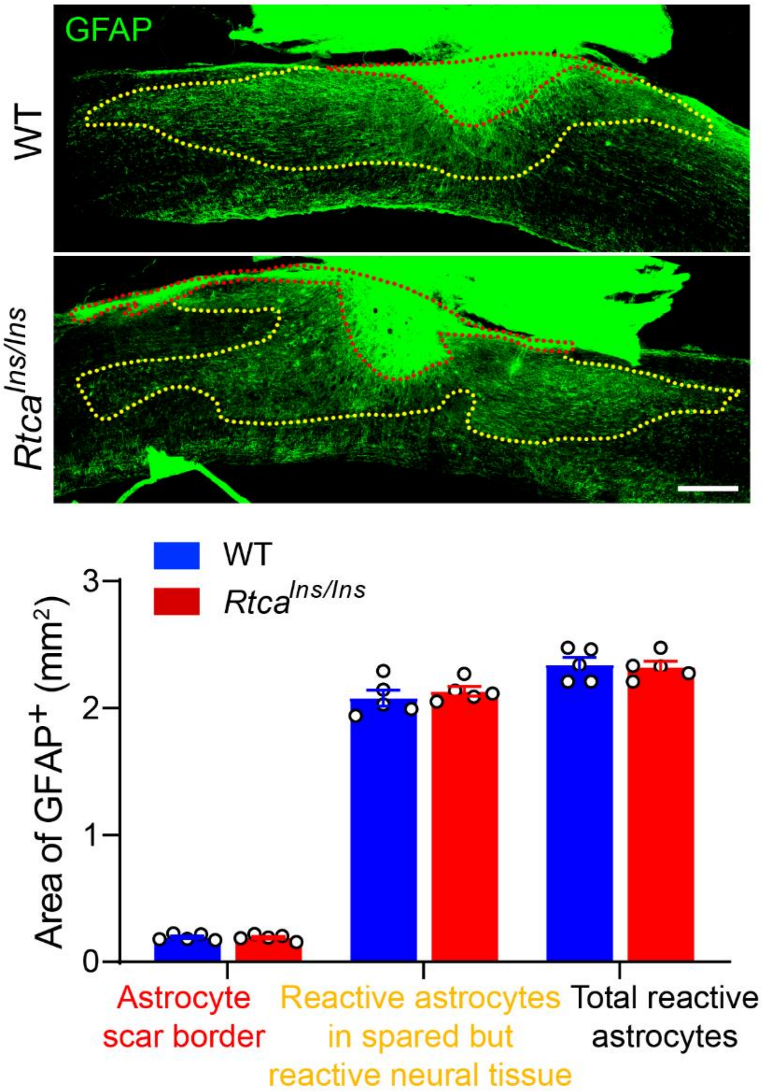


**S2 Figure. Loss of Rtca doesn't alter the glial scar size, related to Figure 2.**

Astrocyte scar border is marked by red dotted line and reactive astrocytes in spared but reactive neural tissue is outlined by yellow dotted line. Analyzed by unpaired t test, $p$ = 0.8380, 0.4946, 0.8355, $n$ = 5. Scale bar, 200 μm.

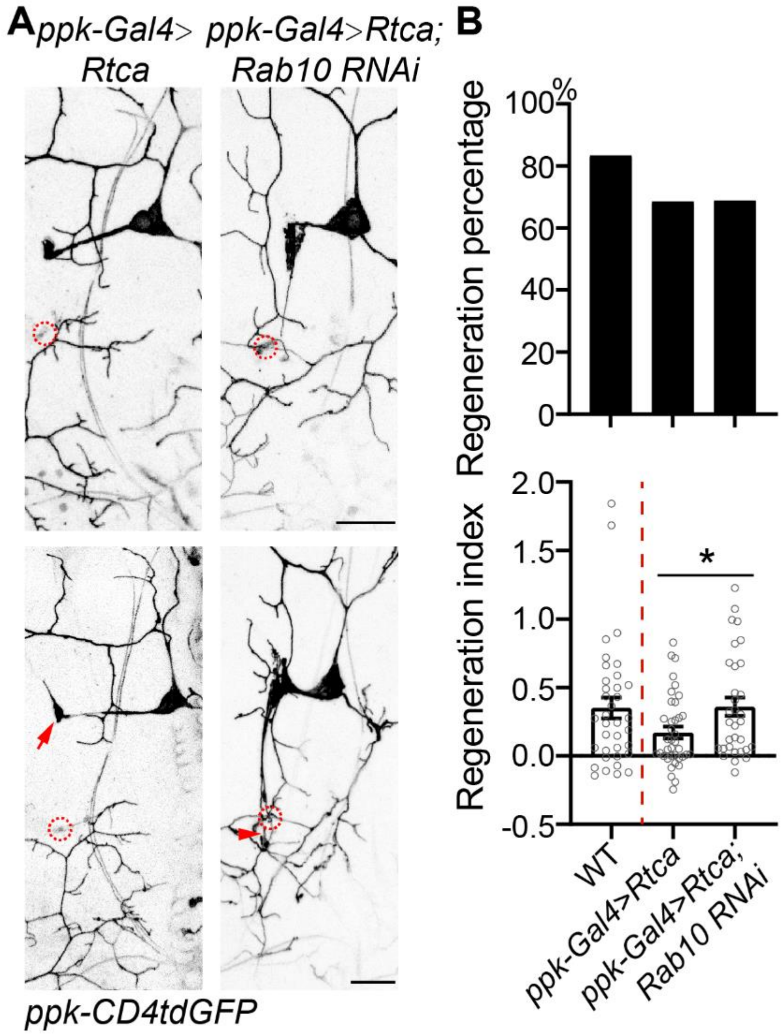


**S3 Figure. Rab10 knockdown rescues the impaired regeneration in Rtca overexpression neurons, related to Figure 4.**

(A) C4da neuron axons were injured and assayed at 48 h AI. The injury site is marked by the dashed circle, regenerating and non-regenerating axons are marked by arrowheads and arrow.

(B) Quantification of axon regeneration by regeneration percentage (upper panel, Fisher's exact test, $p$ = 0.1783, 0.2517) and regeneration index (lower panel, unpaired $t$-test between *ppk-Gal4>Rtca* and *ppk-Gal4>Rtca; Rab10 RNAi*, $p$ = 0.0401). $n$ = 36, 38, 32 neurons. Scale bar, 20 μm. *$p < 0.05$.

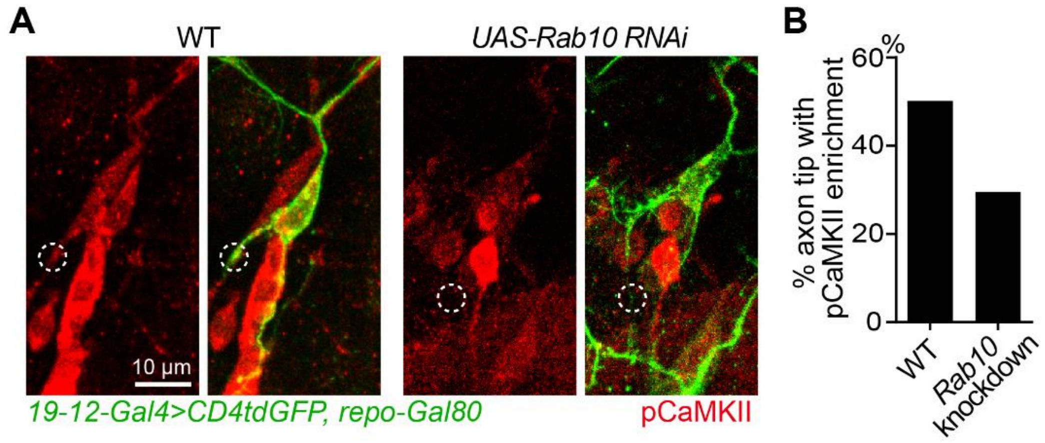


**S4 Figure. Loss of Rab10 impairs the activation of Piezo's downstream component, related to Figure 5.**

(A and B) Piezo's downstream pCaMKII accumulated in 50% WT and 29.41% Rab10 knockdown axon tips at 48 h AI. $n$ = 34 neurons. The axon terminal is marked by the dashed circle. Scale bar, 10 μm.

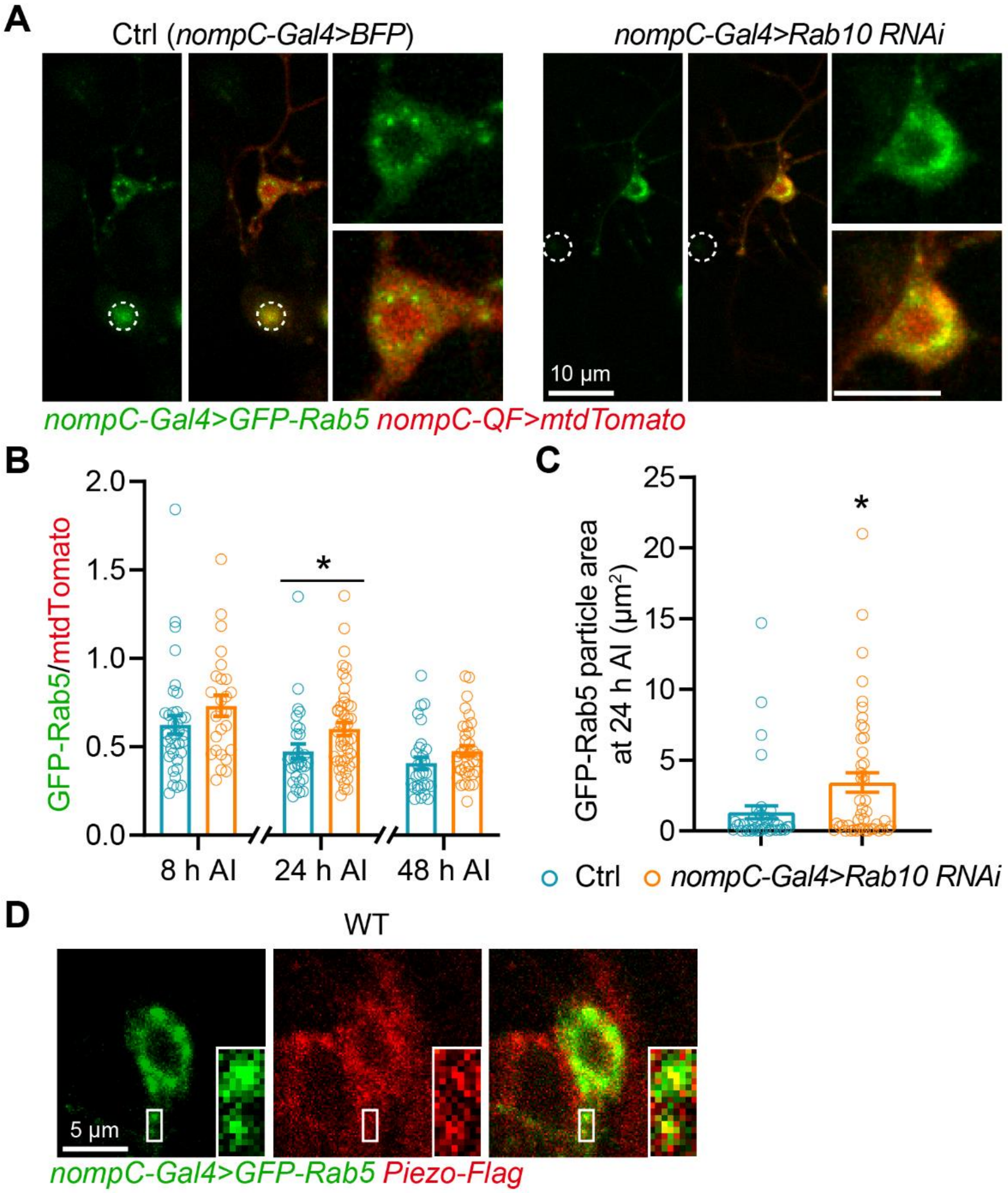


**S5 Figure. Loss of Rab10 affects endosome trafficking, related to Figure 5.**

(A-C) Live image showing GFP-Rab5 fluorescence (B) and GFP-Rab5 positive particle area (C) are increased in Rab10 knockdown neurons at 24 h AI. The injury site is outlined by the dashed circles. Scale bar, 10 μm. (B) Unpaired *t*-test is used to analyze the difference between Ctrl and Rab10 knockdown C3da neurons at 8 h, 24 h, and 48 h AI. $n$ = 36, 26, 30, 47, 29, 35, $p$ = 0.179,

0.0265, 0.124. (C) To assess GFP-Rab5 particle area, the neuron soma was outlined in ImageJ and Analyze Particles was used. Analyzed by unpaired *t*-test, $n = 40, 46$, $p = 0.0153$.

(D) Piezo is detected in GFP-Rab5 positive particles. *Piezo-Flag* knock-in fly stock was generated by fusing Flag to the C-terminal of Piezo. Larvae were dissected at 24 h AI, and their body walls were fixed and stained for GFP and Flag. Scale bar, 5 μm. *$p < 0.05$.

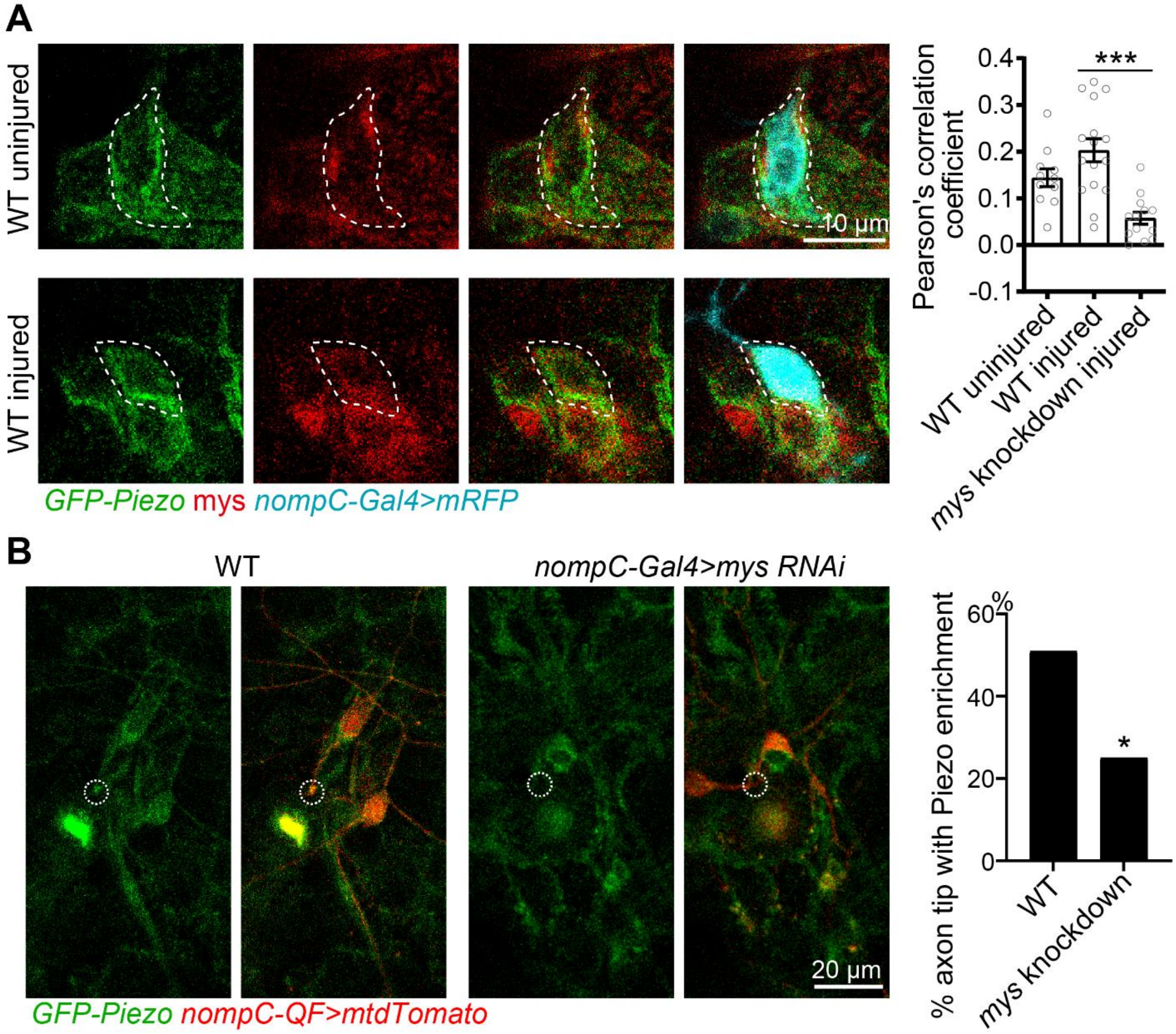


**S6 Figure. mys is required for Piezo membrane targeting, related to Figure 6.**

(A) Piezo's fluorescence signal partially overlaps with mys on the membrane of C3da neurons, and their colocalization modestly increase after injury. To quantify the colocalization between Piezo and mys, the neuron was outlined in ImageJ and assessed with the Colocalization Finder plugin. mys knockdown neuron is served as a negative control. Analyzed by one-way ANOVA followed by Dunnett's multiple comparisons test. $n$ = 11, 16 and 13 neurons. Scale bar, 10 µm.

(B) Piezo's enrichment in the injured axon tip is attenuated in mys knockdown neurons at 24 h AI. The growth cone is highlighted by the dotted white circle. $n$ = 51 and 40 neurons. Scale bar, 20 μm. Analyzed by Fisher's exact test, $p = 0.0172$. *$p < 0.05$, ***$p < 0.001$.

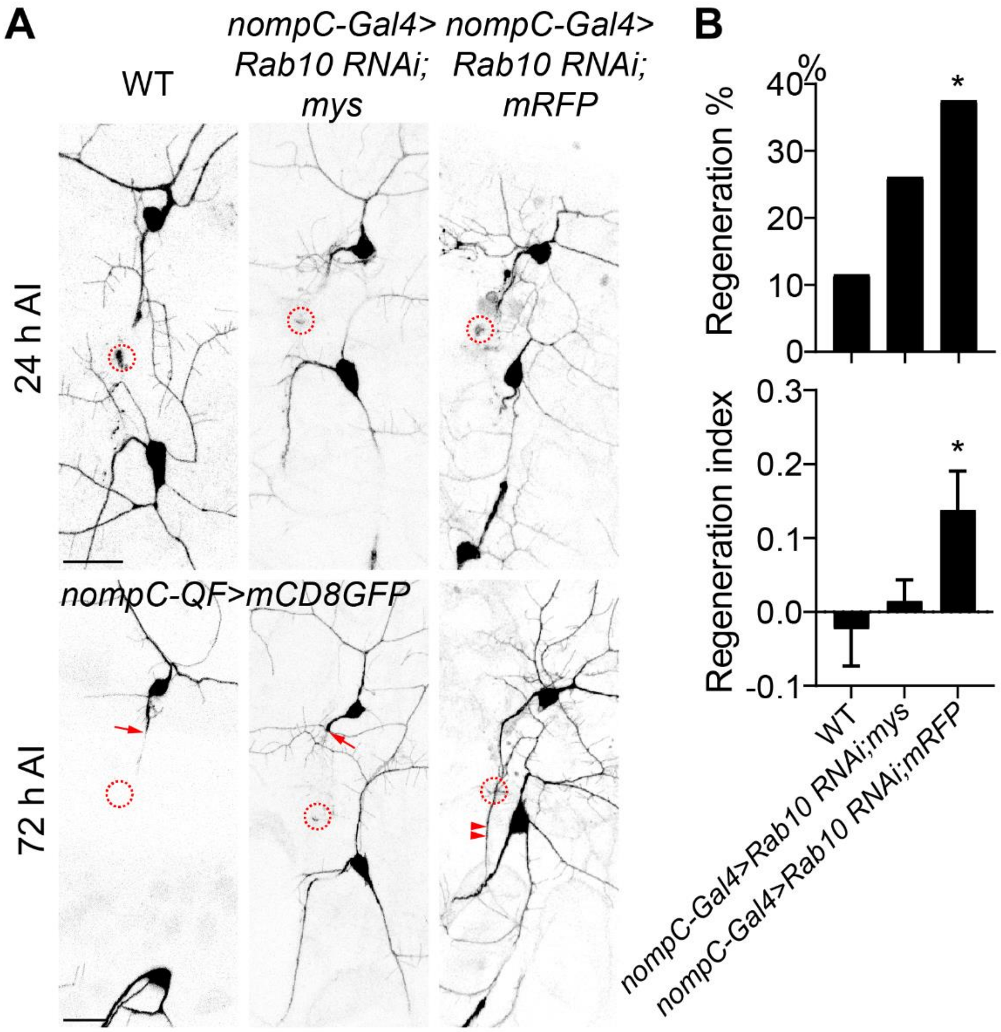


**S7 Figure. Expressing mys attenuates the enhanced regeneration capacity in Rab10 knockdown neurons, related to Figure 6.**

(A) C3da neuron axons were injured and assayed at 72 h AI. The injury site is marked by the dashed circle, the regenerating axon is labeled by arrowheads and non-regenerated axon is marked by arrow.

(B) Quantification of axon regeneration by regeneration percentage (upper panel, Fisher's exact test, $p = 0.2273$, $p = 0.0252$) and regeneration index (lower panel, one-way ANOVA followed by Dunnett's multiple comparisons test). $n$ = 26, 46 and 40 neurons. Scale bar, 20 μm. *$p < 0.05$.

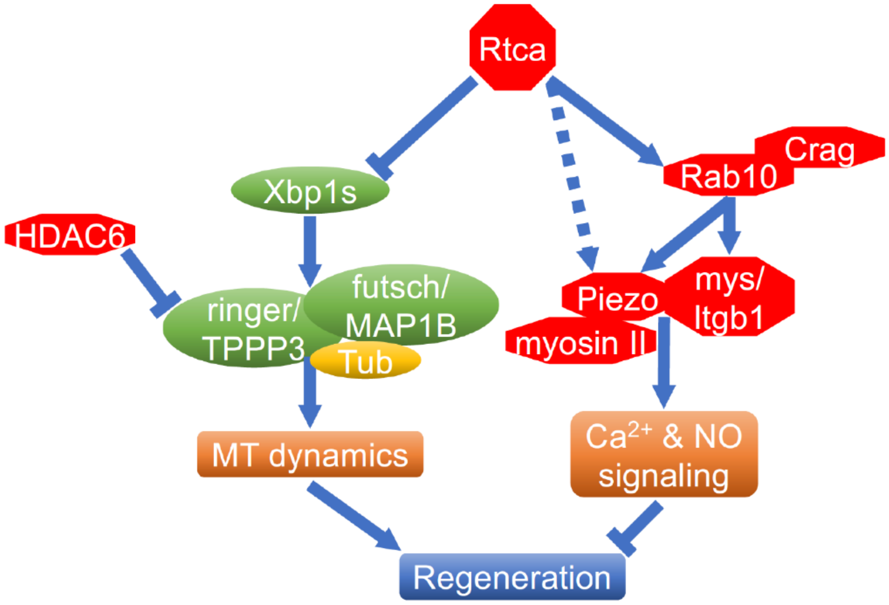


**S8 Figure. Proposed genetic model of the Rtca pathway in mediating axon regeneration, related to Figure 7.**
